# Integrating RNA Structure and Protein Interactions to Uncover the Mechanisms of Viral and Cellular IRES Function

**DOI:** 10.1101/2025.08.29.673120

**Authors:** Riccardo Delli Ponti, Andrea Vandelli, Laura Broglia, Gian Gaetano Tartaglia

**Author notes:** These authors contributed equally.

## Abstract

**Background:** RNAs fold into complex structures that critically influence gene expression. A prominent class of regulatory elements resides in the 5′ untranslated region (5′ UTR), where internal ribosome entry sites (IRESs) promote cap-independent translation by directly engaging the ribosome. First discovered in viral genomes, IRESs have been classified into four types according to their structural compactness and factor requirements. While viral IRESs are well studied, cellular IRESs remain poorly understood: they display limited sequence conservation, reduced structural compactness, and variable dependence on auxiliary RNA-binding proteins known as IRES trans-acting factors (ITAFs). Whether their activity relies mainly on RNA structure or protein assistance remains unresolved. Here, we present a computational framework that combines *in silico* mutagenesis and RNA–protein interaction profiling to investigate IRES mechanisms and guide the design of synthetic elements.

**Results:** Using the Hepatitis C Virus (HCV) IRES as a benchmark, we performed systematic single-nucleotide mutagenesis coupled with structural predictions. Mutations were classified as synonymous or non-synonymous based on their effect on the secondary structure. The HCV IRES showed overall robustness, but the domain interacting with eIF3 was particularly sensitive, consistent with its essential role in translation initiation. Extending this approach to other viral IRES families revealed distinct profiles of resilience: Aphthoviruses retained structural integrity despite extensive sequence variation, whereas Cripaviruses displayed higher variability. We then applied the same analysis to cellular IRESs, which proved more structurally sensitive, suggesting stronger reliance on cofactor support. To probe this connection, we used the *cat*RAPID approach to predict interactions with translation-related proteins. The method distinguished IRESs with known ITAF binding, such as PTBP1, and highlighted stability-promoting mutations that increased the predicted affinity for translation factors.

**Conclusions:** Our *in silico* analysis indicates that mutational tolerance mirrors IRES cofactor dependence: compact viral IRESs are structurally robust, whereas non-viral IRESs are more reliant on protein interactions. By linking structure prediction with interaction profiling, we identify variants that both stabilize IRESs and improve binding to ITAFs or translation factors. This framework provides mechanistic insight into sequence–structure–function relationships and supports the rational design of synthetic IRES elements for therapeutic and biotechnological applications.

## Background

RNA molecules can fold into complex structures and adopt specific three-dimensional conformations. Their structural complexity and flexibility enable diverse functions—from catalytic activity in ribozymes to environmental sensing by metabolite-responsive riboswitches. One region where RNA structural elements converge is the 5’ untranslated region (5’ UTR), which can host a diverse array of functional RNA structures, including internal ribosome entry sites (IRESs) (1). IRESs were first identified in viral genomes, where they sustain translation by hijacking the host’s protein synthesis machinery. These structured elements enable the direct recruitment of the 40S ribosomal subunit, bypassing the need for cap-dependent initiation—an advantage during viral infection when host cap-dependent translation is inhibited (2–5). Despite a shared functional role in promoting cap-independent translation, viral IRESs display remarkable diversity in their nucleotide sequences, secondary structures, and reliance on host cofactors. These proteins include the eukaryotic initiation factors (eIFs) and auxiliary RNA-binding proteins (RBPs), known as IRES *trans*-acting factors (ITAFs). Based on these features, viral IRESs are classified into four major types according to their requirements for ribosome recruitment: Type I (*e.g*., Poliovirus, PV) and Type II (*e.g.*, Foot-and-Mouth Disease Virus, FMDV) IRESs depend on multiple canonical eIFs (excluding eIF4E) and ITAFs (6,7); Type III IRESs, exemplified by the Hepatitis C Virus (HCV), require exclusively the eIF3 and the eIF2 ternary complex. The HCV IRES engages in direct base pairing with the 18S rRNA, inducing a conformational change in the 40S subunit and facilitating its recruitment. In particular, the interaction between eIF3 and a specific stem-loop in the IRES modulates the efficiency of pre-initiation complex assembly (8). In contrast, Type IV IRESs, such as that of Cricket paralysis virus (CrPV), operate entirely independently of host factors, directly recruiting the ribosome by structurally mimicking tRNA–mRNA interactions (9,10). Overall, the more structured and compact an IRES is, the less it relies on host cofactors for ribosome recruitment. In contrast, IRESs with less intrinsic structure tend to depend more heavily on eIFs and ITAFs to achieve translation initiation (9).

While viral IRESs have been extensively studied, non-viral IRESs remain poorly understood. Several non-viral mRNAs, particularly those involved in stress responses or encoding transcription factors, contain IRES elements in their 5’ UTRs (11). Although cap-dependent translation dominates under homeostatic conditions, non-viral IRESs can drive gene-specific translation, especially during stress or development. Genome-wide studies suggest that a notable fraction of human 5’ UTRs may harbor IRES-like activity (12), yet the structural and mechanistic basis of this function remains elusive.

In contrast to viral IRESs, cellular IRESs show low sequence conservation, reduced structural compactness, and limited similarity to one another (13,14). These elements are often highly dynamic, undergoing conformational changes that modulate their activity (15–17). Only a handful of non-viral IRESs have been characterized in depth. Notable examples occur in the mRNAs of key regulators of proliferation and differentiation—such as *c-MYC* (*18,19*), *VEGFA* (20–22) and *IGF2* (*23,24*). These IRESs rely on the activity of ITAF such as PTB1 for *c-MYC* IRES (19) and *cis-*acting elements as for instance upstream open reading frames (uORFs) and G quadruplex (RG4) as in the case of *VEGFA* (25–27). While some ITAFs are shared between viral and non-viral IRESs, most remain unidentified, and their precise role—whether in IRES stabilization or structural remodeling—remains unclear (28,29).

Although the secondary and tertiary structures of viral IRESs are relatively well characterized, their direct interactions with the ribosome and ITAFs are still incompletely defined. This knowledge gap is even more pronounced for cellular IRESs, where both the contribution of RNA structure and the identity of regulatory RBPs remain largely unresolved. Here, we integrated RNA structure prediction and RNA–protein interaction profiling to explore how sequence variation affects RNA folding, interaction propensity with proteins, and thus IRES function. Using the HCV IRES as a model, we classified single-nucleotide variants based on their predicted structural impact and identified regions of differential sensitivity to mutations. We then extended this analysis to other viral and non-viral IRESs, revealing distinct mutational tolerance across viral and non-viral IRESs. Mutational tolerance is element-specific and largely architecture-dependent, ranging from broad susceptibility to single-nucleotide changes (*e.g.* CrPV) to sensitivity confined to discrete structural modules (*e.g.* HCV and FMDV). We then applied *cat*RAPID (30,31), a method to predict protein-RNA interactions, to evaluate changes in ITAFs and eIFs binding. By linking the structural consequences of mutations with predicted changes in protein-binding affinity, we aim to infer functional relevance from the mutational profile. This enabled the identification of mutations that are likely to enhance IRES activity—by stabilizing favorable structures and by promoting interactions with key components of the translation machinery. Altogether, this approach offers a framework not only for understanding how sequence and structure influence IRES function, but also for the rational design of synthetic IRES elements with improved translational activity.

## Methods

### RNAfold and Cryo-EM structures

We extracted the sequences for domain I and III of HCV IRES directly from the paper of Quade et al, specifically from Figure 1A (3). After selecting the sequences, we used the webserver version of RNAfold to predict their MFE structure (32). We then compared the dots-and-brackets native and predicted structures, and used the positive predicted value (PPV; number of matching brackets) and the structural identity (number of matching dots and brackets) to evaluate the predictions.

**Figure 1.**
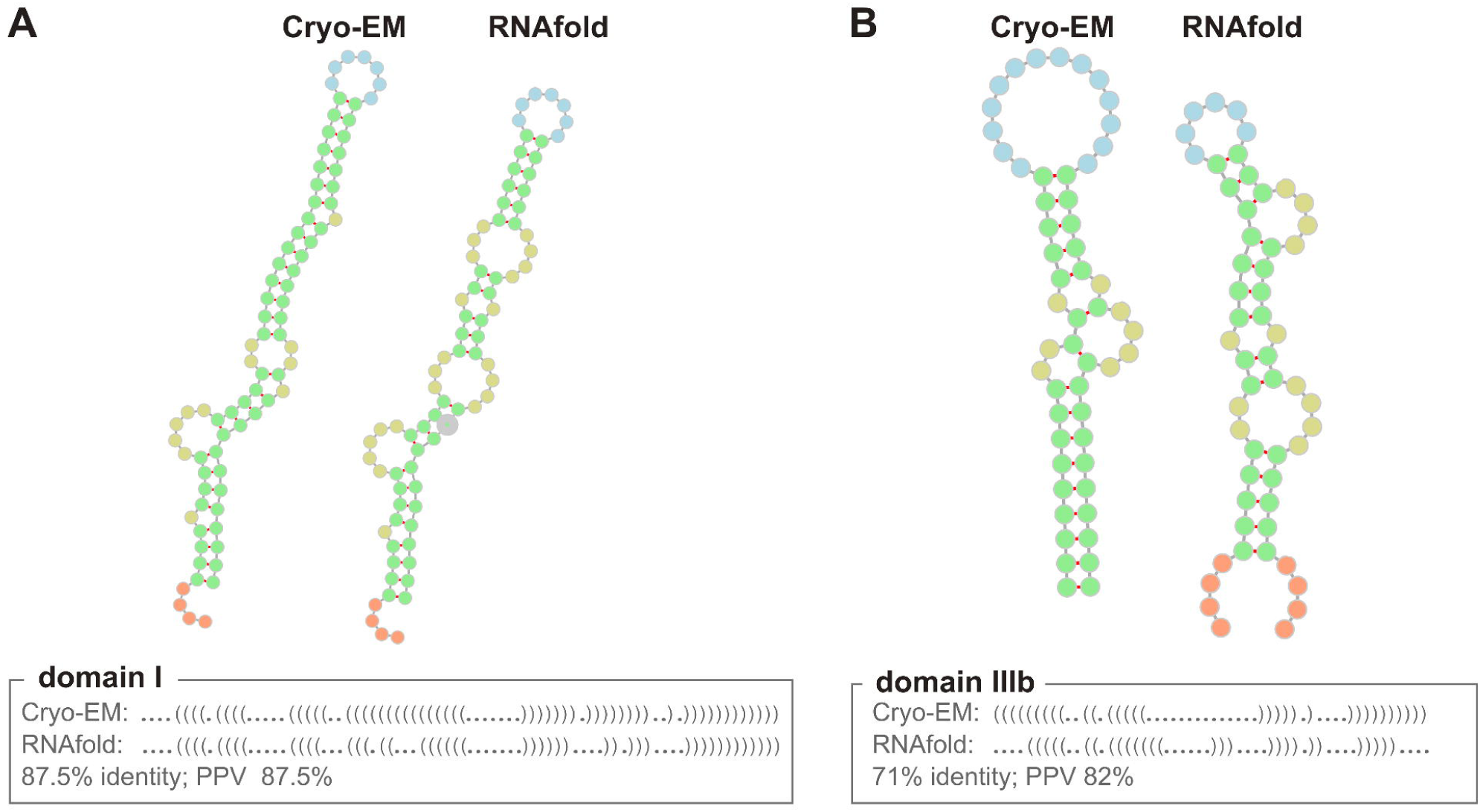
Comparison between predicted and experimentally determined HCV structures. Comparison of Cryo-EM secondary structure and RNAfold prediction for the domain I (**A**) and IIIb (**B**). The images were obtained using the FORNA web app. The nucleotides are colored by the web app according to their structure, for example stems in green and loops in blue (114).

### Building the IRES dataset

We selected the IRES sequences available on the RFAM website (33): HCV (RF00061), Apthoviruses (RF00210), Picornaviruses (RF00229), Cripaviruses (RF00458), *c-MYC* (RF00216), *VEGF* (RF00461), *IGF2* (RF00483). For each IRES family, we downloaded all the available sequences and used them as our starting dataset. From Picornaviruses, we only extracted the Poliovirus sequences, and for Apthoviruses, only Foot-and-Mouth Disease virus.

### Mutation profiling

First, we scanned the available sequences, one nucleotide at a time. Then, the selected nucleotide was randomly mutated into another nucleotide (A→C, G, U; C→A, G, U; G→C, A, U; U→A, C, G). The resulting single-nucleotide mutant was then used as the input sequence for RNAfold, launched from the command line using default values. We then compared the structure of each mutant with the wild-type structure, using the structural identity to evaluate the models.

### Selecting structural beneficial mutations

For each mutant structure, we also computed the free energy (RNAfold output) and the structural content (i.e., % of double-stranded nucleotides). The information coming from the structural identity with the native sequence, the difference in free energies, and the difference in the structural content was employed to select what we defined as “structurally beneficial” or stability-enhancing mutations. Specifically, we selected these mutations according to: (i) structural identity ≥ 90% relative to the wild-type; (ii) increased structural content (SC), measured as a higher percentage of base-paired nucleotides (SC Mut > SC WT); (iii) lower predicted minimum free energy (ΔG Mut < ΔG WT).

### Protein–RNA Interaction Predictions

Interactions between IRES sequences and human RNA-binding proteins (RBPs) were predicted using *cat*RAPID, an algorithm that estimates the binding propensity of protein–RNA pairs. The model integrates multiple features, including RNA secondary structure, hydrogen bonding, and van der Waals forces. It was trained on approximately half a million experimentally validated interactions and has demonstrated robust performance in distinguishing interacting from non-interacting pairs (AUC = 0.78) (31,34).

For PTBP1 and selected ITAFs, we used the *cat*RAPID interaction propensity scores of wild-type (WT) IRES sequences to assess binding strength and compare results across different sequences.

### Interactions with human translation-related RBPs

To identify the most promising candidates among beneficial IRES mutations, we predicted their interactions with a curated dataset of 119 human RBPs extracted from the UniProt database (Release 2025_03) (35), applying the following filters:

- Reviewed status
- Annotated with translation-related functions (GO:0006412)
- RNA-binding activity (GO:0003723)
- Excluding ribosomal proteins (GO:0044391)
- Excluding translation repressors (GO:0030371)

This approach allowed us to focus on RBPs that generally enhance translation but are not part of the ribosomal machinery or translational repressors.

For each mutated sequence, we calculated the percentage of proteins binding to it for which the mutation resulted in an increase in interaction propensity >10, identifying those mutations most likely to enhance translation via stronger RBP binding.

### Analysis of *c-MYC* IRES Mutants

To validate the potential regulatory effects of mutationA and mutationD within the *c-MYC* IRES, we predicted their interactions against a larger panel of 2,064 canonical and non-canonical human RBPs (31). For each protein, we computed the difference in *cat*RAPID score between the mutated and WT sequences, prioritizing those whose binding was enhanced by the mutation. The top 50 RBPs with the highest positive deltas were selected and subjected to Gene Ontology (GO) enrichment analysis using the tool g:Profiler (36).

### *VEGF* and *IGF2* IRES Mutant Screening

We also examined the binding propensity between mutated versions of the *VEGF* and *IGF2* IRESs and the same set of 2,064 RBPs. Only mutations showing a binding propensity improvement >10 (compared to the WT) were retained for each RBP. For each protein, we averaged the differences from all these beneficial mutations to compute a summary score and rank the proteins accordingly. The top 100 RBPs were then analyzed for GO enrichment using g:Profiler (36).

### Interactions with human ITAFs

We predicted the binding propensity between WT sequences of *c-MYC*, *VEGF* and *IGF2* IRESs with the set of 2,064 RBPs. We ranked the proteins according to their average binding propensity. For *c-MYC*, for different percentages of ranked RBPs, we calculated the enrichment of known human ITAFs (37,38) compared to 10 random interactomes. The enrichment is calculated as a Z-score for each percentage with average and standard deviation derived from both target and random interactomes.

### Sequence conservation of ITAFs

The conservation of ITAFs across different organisms was retrieved on UniRef clustering within the UniprotKB database (39).

## Results

### Computational Analysis of HCV IRES Structural Robustness and Mutational Impact

Given its well-characterized structure and the extensive availability of experimental data, the HCV IRES was selected as a reference model for our computational analyses (3,40–44). The HCV IRES exemplifies a type III IRES element, which, while capable of directly interacting with the 40S ribosomal subunit, is not fully independent (43,45,46). As the first step, we validated our structural predictive approach by using RNAfold (32) to model the IRES structure and compared it with its cryo-EM structure (3). RNAfold achieved good performances against the domain I (40-119 nt; Identity= 87%; PPV= 87%) and IIIb (173-227 nt; Identity= 71%; PPV= 82%) of HCV IRES (see **Material and Methods**) (**Figure 1**). Given the accuracy of RNAfold in modeling HCV IRES, we extended its use to investigate the structural robustness of this element under mutational pressure, with the aim of uncovering the structural basis of its function.

In all our analyses, IRES sequences were retrieved from the RFAM database (33) (see **Material and Methods**). In particular for HCV, we selected all the sequences of the RF00061 family, for a total of 52 sequences. For every sequence of the HCV IRES dataset, we randomly mutated every single nucleotide and then predicted the secondary structure of each resulting variant (see **Material and Methods**). The structure of every mutant (MT) was then compared with the wild-type (WT) to assess the effect of mutations on the IRES structure. Mutants with a structural identity of 100% (which we call hereafter “**synonymous structural mutations**”) retain the same structure of the WT, indicating that these mutations do not disrupt RNA folding. On the contrary, mutants showing 50% sequence identity with the WT (*i.e*., “**non-synonymous structural mutations**”) exhibit single-nucleotide changes that markedly alter the overall IRES secondary structure (**Figure 2A**). First, we did not observe striking differences based on the specific nucleotide substitutions, although transversions mutations such as G to C and C or U to G showed a slightly greater impact on structural disruption (**Supplementary Figure 1**). This suggests that the impact of a mutation on the overall IRES structure—and consequently its function—is more dependent on the position of the substitution than on the specific nucleotide change, indicating a predominantly position-dependent rather than base-specific effect. Moreover, previous mutation profiles performed for HCV stem loop IV also showed that certain point mutations preserve the original structure highlighting intrinsic structural robustness. Their analysis further revealed that base-paired positions evolve simultaneously and that not all nucleotide positions contribute equally to structural stability—some being evolutionarily flexible, while others are energetically constrained (47).

**Figure 2.**
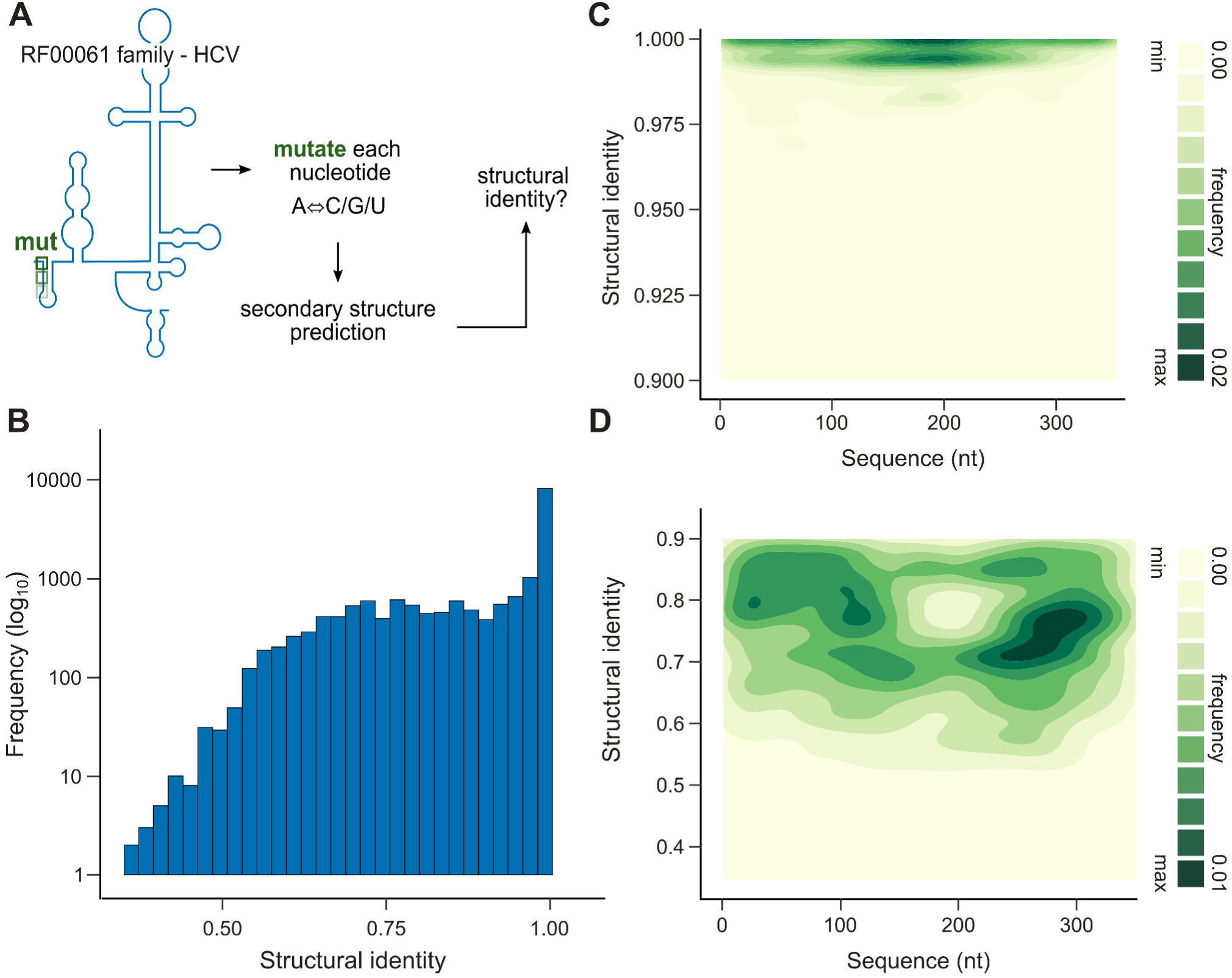
Mutation distribution and structural identity in HCV IRES variants. **A.** Schematic representation of the single-nucleotide mutagenesis of the HCV IRES (RF00061). For each variant we predict the fold and compute structural identity relative to the WT. Green marks indicate example mutations. **B.** Number of mutations and the related structural identity when compared with the WT IRES sequences **C.** Density of mutations in HCV IRES sequences with structural identity >U0.90. Mutation density is color-coded from low (light yellow) to high (dark green). **D.** Density of mutations with a structural identity <0.90 in HCV IRES sequences. The density of mutation goes from low (light yellow) to high (dark green).

The majority of the mutations do not alter the RNA secondary structure (**Figure 2B**), with 23% of the total mutations showing 100% structural identity. This pattern is consistent with genome-scale analyses showing that RNA secondary structure constrains sequence variability in RNA viruses (48–53). However, we also identified a small subset of mutations (∼0.5%) that cause a significant disruption of the IRES secondary structure (structural identity < 50%). Interestingly, some synonymous structural mutations were found to stabilize (lower free energy) or destabilize (higher free energy) the overall structure. In the majority of the cases, they do not affect the energy (**Supplementary Figure 2**). When mapping these mutations onto the HCV IRES, we observe that most substitutions preserving the secondary structure (structural identity > 90%) cluster predominantly within the central region (nucleotides 150–175; **Figure 2C**). In contrast, mutations that substantially disrupt the secondary structure (structural identity < 90%) are more frequently localized toward the 3′ region of the IRES (**Figure 2D).**

We next investigated the mutations altering the IRES secondary structure with different intensity (structural identity <70%, <60%, <50%; **Figure 3**). Interestingly, mutations slightly altering the secondary structure (structural identity < 70%) are more prominent in the 3′ region of HCV IRES. On the other hand, the mutations altering the structure (structural identity < 50%) are more concentrated in the region at nucleotides ∼150-250, which is part of the IIIb domain responsible for binding to eIF3, especially nucleotides 175-200. This result suggests that the eIF3 binding region is more susceptible to mutations. It must be noted that HCV-like IRESs interact with eIF3 and this interaction is crucial for the IRES-mediated translation initiation (8,43,45,54–57). The binding of eIF3 to the apical region of HCV domain III serves a dual role: it reduces competition for the 40S ribosomal subunit and limits the formation of 43S pre-initiation complexes, thereby hindering the translation of non-viral mRNAs thus promoting the viral RNAs (54). Mutations within the eIF3-binding region have been shown to impair IRES-mediated translation in cell-free extract assays (58). Because this domain retains a conserved stem–loop structure across HCV isolates despite sequence variation, it indicates that eIF3 recognizes RNA structure rather than the primary sequence (59,60). Overall, these observations align with our results, which indicate that the eIF3-interacting region is particularly sensitive to mutations that disrupt its secondary structure.

**Figure 3.**
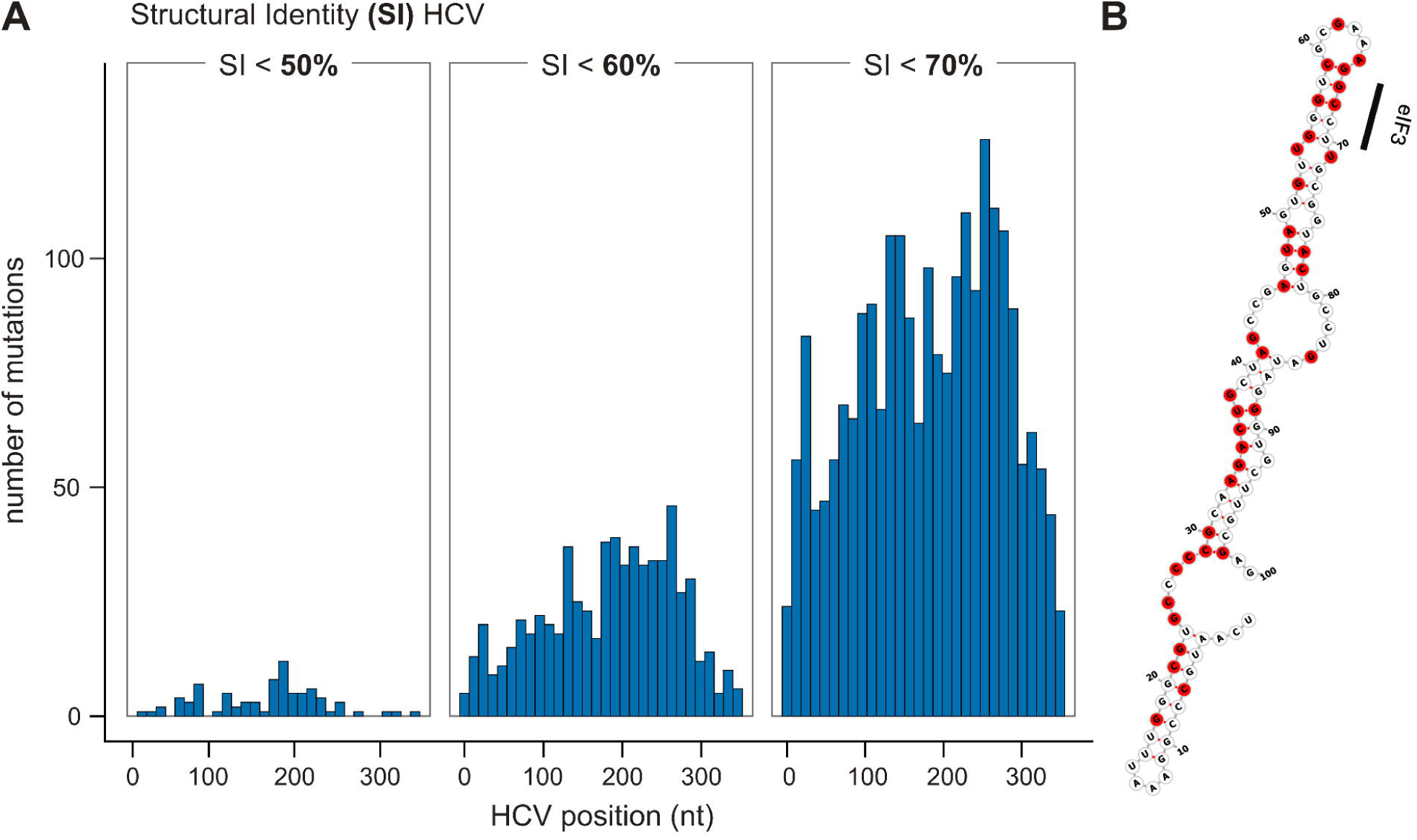
Mutation profiling of HCV IRES. **A.** Position on the HCV IRES of the mutations affecting the structure, with structural identity (SI) < 50%, 60% and 70%. **B.** Secondary structure of HCV region 150-250nt, the region with highest presence of mutations affecting the structure (structural identity < 0.5). Nucleotides are colored according to their minimum structural identity (SI) of the mutation identified, specifically red (SI < 0.5).

The presence of mutations that drastically disrupt the IRES secondary structure—and likely impair its function—also implies the potential existence of mutations that enhance IRES activity. Deletion of domain I of the HCV genome has been shown to increase translation activity (58,61), suggesting that this region may function as an inhibitory element of the IRES. Not only large deletions but also single-nucleotide changes can enhance IRES function. For example, nucleotide insertions at the 3′ end of domain IIIb have been reported to boost IRES activity (60,62), likely by increasing the affinity for eIF3 binding and thereby promoting more efficient translation initiation. Moreover, U→C substitutions that disrupt base pairs within the P3.1 helix of the CrPV IGR IRES increase IRES activity by facilitating ribosome loading. Likewise, the A6187G substitution in PKI creates a G·U wobble enhances activity, likely by favoring the pseudo-translocation step (63). The same is true for non-viral IRES, for instance, a single nucleotide mutation (C→U) within the human c-MYC IRES has been shown to enhance its activity (64), likely by promoting the formation of a stem-loop structure and favoring the recruitment of the ITAFs PTB1 and YB-1 (19,64,65).

### Synonymous and non-synonymous structural mutations in viral and cellular IRESs

We selected representative viral IRESs from each of the four established classes, which are defined based on their structural organization and requirements for translation initiation factors: type I (*Enterovirus, e.g.* PV), type II (*Aphthovirus*, *e.g.* FMDV), type III (*Hepacivirus*, *e.g.* HCV) and type IV (*Cripavirus*, *e.g* CrPV). For each viral IRES, we generated the mutation profiling and compared it with the WT, following the workflow applied to HCV IRES. While the viral IRESs retain the majority of their structural identity, their tolerance to single-nucleotide mutations differs across different IRES types (**Supplementary Figure 3**). Aphthoviruses, specifically FMDV, demonstrate high structural resilience, retaining much of their native secondary structure upon single-nucleotide mutations. In contrast, Cripaviruses exhibit greater structural variability following mutational perturbation (**Supplementary Figure 3A**), meaning that larger fraction of substitutions substantially reconfigure base-pairing (lower robustness). Type IV IRESs consists of a quite compact and rigid structure which is essential to promote factor-independent association to the ribosome (66). Therefore, disrupting base-pairing could impair function, indicating limited tolerance to sequence changes (67,68).

To relate structural tolerance to regulatory context, we also included non-viral IRESs in the analysis (**Supplementary Figure 3B**). Specifically, we selected from RFAM the IRES of *c-MYC*, *VEGF*, and *IGF2* (**Materials** and **Methods**). Non-viral IRESs differ markedly from their viral counterparts, both in evolutionary origin and functional context. Viral IRESs are essential for efficient translation and viral fitness, and thus their structures are under constraints to ensure conservation despite high mutation rates (69). In contrast, mammalian IRESs are highly regulated by the cell and are often activated under specific stress conditions and environmental changes that suppress cap-dependent translation (14,70). Non-viral IRESs are less structured compared to viral IRESs (14,38), and they depend heavily on eIFs and ITAFs to fold or remodel into a translation-competent conformation. Accordingly, our single-nucleotide-mutation profiling shows that the *c-MYC* and *VEGF* IRES elements behave much like type I/II viral IRESs; these non-viral elements contain discrete RNA segments—probably ITAF-binding sites—that must remain intact for proper function. Notably, among the IRESs analyzed, the *IGF2* IRES displays the highest degree of structural variability following mutations (**Supplementary Figure 3**). Unlike the more compact *c-MYC* and *VEGF* IRESs, *IGF2* adopts a loose fold of four small stem–loops and lacks several structural motifs common to other non-viral and viral IRESs (24).

To further investigate the effects of single-nucleotide mutations on the secondary structure of viral and non-viral IRES elements, we focused on changes that severely disrupt folding (structural identity < 50%; **Figure 4**). As observed previously, viral IRESs generally exhibit a lower frequency of such disruptive non-synonymous structural mutations (**Figure 4A**). The CrPV IRES displayed the greatest mutational susceptibility, with numerous positions where single point mutations substantially alter the predicted secondary structure especially when compared to FMDV which exhibits a compact, 3′-terminal region with concentrated sensitivity, falling within the highly structured and conserved domain III of the IRES (71).

**Figure 4.**
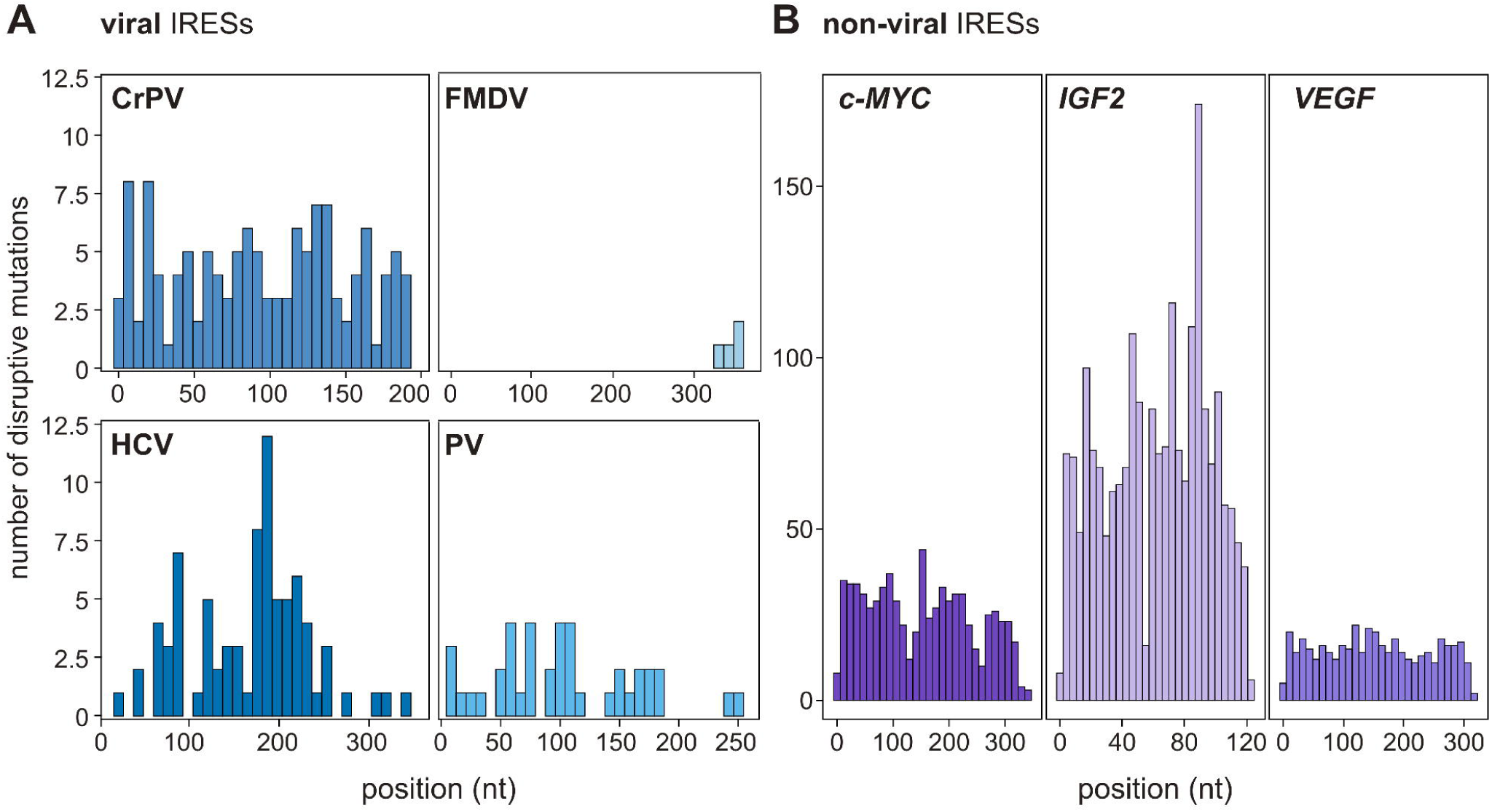
Distribution of structure-disruptive mutations across viral and non-viral IRESs. The position of the mutations highly affects the secondary structure (structural identity < 50%) on the viral (**A**) and non-viral (**B**) IRES sequences.

When we applied the same analysis to non-viral IRESs (**Figure 4B**), we observed a higher incidence of disruptive single-nucleotide changes and more extended segments of sensitivity than in most viral elements (**Figure 4A**) This observation reinforces the idea that non-viral IRESs operate under less stringent constraints than viral IRESs. The structure–function relationship of non-viral IRESs is comparatively flexible compared to viral IRESs (72). Their activity is buffered by an extensive regulatory network that includes trans-acting factors— helicases, RNA-modifying enzymes, and nuclear-cytoplasmic shuttling RBPs—and cis-acting elements such as upstream open reading frames (uORFs) and G-quadruplexes, together allowing for plasticity without complete loss of function (1,73). In summary, the discovery of mutations that dramatically remodel IRES architecture—and, in turn, its function—suggests that few sequence changes are not tolerated but may actually enhance IRES-driven translation.

### IRES and interaction with ITAFs

Before investigating whether certain mutations enhance IRES activity, we first examined the set of proteins that bind the IRESs. We reasoned that strengthening the interaction with positive regulators—such as specific ITAFs or translation initiation factors—could represent a mechanism by which some mutations indirectly promote IRES function. IRES elements, both cellular and viral, often rely on the assistance of RBPs (ITAFs) and components of the translation machinery, such as eukaryotic initiation factors (eIFs), to function efficiently (14,74). These regulatory proteins are frequently shared between non-viral and viral IRESs, highlighting a common dependence on host factors to support cap-independent translation. We used the *cat*RAPID algorithm to study the protein-IRES interaction *in silico* (*31*). *cat*RAPID was previously successfully employed to study the interaction of functional viral RNA structures and the human proteome, including the XRN1-resistant structures in flaviviruses (75), and human-SARS-CoV-2 interaction networks (76,77).

First, we examined the interaction between the HCV IRES and its essential translation initiation factors, eIF2 and eIF3 (43,78–80). Indeed, while the HCV IRES directly contacts the 40S it requires eIF3 and the GTP-eIF2-Met -tRNAi to position the ribosome at the correct AUG site and initiate translation. Using the WT IRES sequences (without any introduced mutations), we employed *cat*RAPID to predict binding affinities with eIF2 and eIF3. We further evaluated the predictive performance of *cat*RAPID by including two control datasets. As a positive control, we retrieved 5′ UTRs of protein-coding RNAs from Ensembl BioMart (81), matched in length to the HCV IRES (approximately 250–350 nucleotides). As a negative control, we selected long non-coding RNAs (lncRNAs) of similar length. We then applied *cat*RAPID to all sequences to compute their interaction propensities with eIF2 and eIF3 (**Figure 5A**). As expected, the 5′ UTRs showed the highest binding scores. Notably, the HCV IRES exhibited stronger binding than the lncRNA controls (eIF3A: p-value < 1.3×10^-8^, EIF2S1: p-value < 2.5×10^-17^), confirming its capacity to robustly interact with the translation machinery, particularly eIF2 and eIF3.

**Figure 5.**
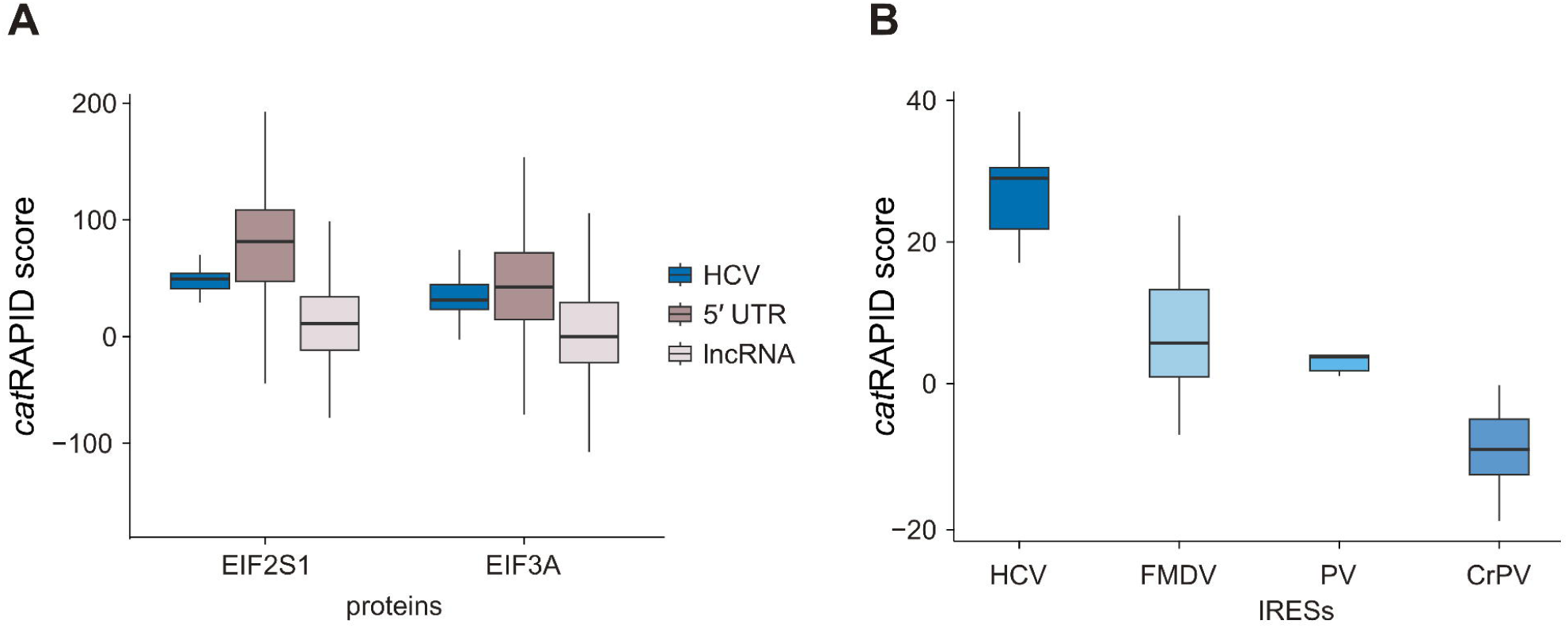
Prediction of IRES interaction propensity with known ITAF. **A.** Predicted binding propensity of HCV IRES with eIF3A and eIF2S1; *cat*RAPID score of WT sequences is reported on the y-axis. **B.** Predicted binding propensity of PTBP1 with different viral IRES sequences. *cat*RAPID score of WT sequences is reported on the y-axis.

In addition, each IRES typically engages a distinct set of regulatory ITAFs, which are not always directly involved in the core translation machinery. Indeed they are often nuclear RBPs that translocate to the cytoplasm in response to specific environmental or stress conditions (74). To facilitate a broader comparison of protein–IRES interactions across diverse viral elements, we selected PTBP1 as a representative “universal” viral ITAF, as multiple binding sites for this protein have been identified in IRESs from different viral families, including PV, EMCV, HCV, and FMDV (74,82).

We used our *in silico* approach to predict the interaction of PTBP1 with all the viral IRES sequences. HCV and FMDV have the highest interaction, with a positive *cat*RAPID score. CrPVs are instead predicted not interacting (*cat*RAPID score < 0 indicative of poor interaction propensity; **Figure 5B**). This result is consistent with previous findings demonstrating PTBP1 binding to the PV (83), HCV (84) and FMDV (85) IRESs, and further highlights the limited interaction between CrPV IRESs and host proteins. This weak host-protein engagement supports the notion that CrPV IRESs can recruit the ribosome independently of protein cofactors. These results support the use of *cat*RAPID to establish *in silico* interaction between IRES sequences and known ITAFs or elements of the translation machinery.

To further assess whether human IRESs show a preference for binding known ITAFs and to demonstrate the effectiveness of our predictive strategy, we collected a curated list of experimentally validated human ITAFs (37,38), and used *cat*RAPID to predict the interactions of *c-MYC*, *VEGF*, and *IGF2* IRES sequences against the dataset of >2,000 human RBPs. For *c-MYC*, we ranked proteins according to their average binding propensity and quantified the enrichment of known ITAFs at different thresholds, comparing the results against 10 random interactomes generated from the same set of RBPs (see **Materials and Methods**). Interestingly, the top 1% of *c-MYC* interactions showed a strong enrichment in known ITAFs (**Supplementary Figure 4A**). This observation underscores the central role of ITAFs in the regulation of human IRESs. We identified YBX1, a previously reported *c-MYC* ITAF, as having the highest binding propensity for *c-MYC* (86). Notably, it also emerged among the top 5% of ranked RBPs for both *VEGF* and *IGF2* (**Supplementary Figure 4B**). Previous studies have demonstrated that YBX1 is critical for the translation of HCMV mRNA (87) and it can bind the wild type p16INK4a mRNA increasing its translation efficiency during hypoxic stress (88). Furthermore, this protein seems to act as a ITAF for HIF1-α mRNA and modulates the expression of VEGF165 and VEGF165b (89), promoting the metastatic capabilities of sarcoma cells. This suggests the key role of YBX1 in IRES-mediated regulation and translation and highlights the strength of our predictive method.

### *c-MYC* IRES as a model for activity-enhancing mutations

While non-synonymous mutations can disrupt RNA folding and compromise IRES function, certain synonymous structural mutations—which preserve the overall fold while subtly modulating local features—can enhance translational activity. This principle is exemplified by a naturally occurring C→U mutation in the c-MYC IRES, discovered in cell lines derived from multiple myeloma patients (64,65). This single-nucleotide change promotes the formation of an additional stem–loop structure near the pseudoknot region, stabilizing the RNA without disrupting its global architecture. Functionally, it leads to enhanced IRES-mediated translation, correlated with increased binding of the ITAFs PTBP1 (19) and YBX1, likely facilitating more efficient ribosome recruitment. These results indicate that sequence– structure–function tuning via minimal mutations can yield functionally beneficial outcomes.

Optimization of IRES activity can improve protein output in synthetic constructs and support applications in vaccine design and regulatory RNA engineering (90–93). Building on this premise, we aimed to provide an exploratory landscape to engineer an optimized IRES *in silico*, using the *c-MYC* IRES as a case study. In the first secondary-structure characterization of the c-MYC IRES, targeted mutations were introduced to probe the contribution of individual structural elements. Notably, two four-nucleotide loop substitutions—MutA (GGGAA→GAAUU) in domain 1 and MutD (AUUU→GAAA) in domain 2—each increased IRES activity when tested individually (90). These mutations affect apical loop structures thought to mediate RNA–protein interactions, and their effects reinforce the importance of fine-scale RNA architecture in regulating translation initiation. In particular, MutD produces a stronger translational enhancement than MutA, making it a more robust reference point for downstream analysis.

To benchmark our IRES-optimization approach, we first analyzed the effects of MutA and MutD. Structurally, both mutations retained the global architecture of the IRES: MutA preserved 97% structural identity, while MutD retained 90%. However, the thermodynamic and structural impacts diverged. MutD lowered the free energy and increased structural content, suggesting a more stable and potentially more protein-accessible structure. In contrast, MutA led to increased free energy and a reduction in base pairing, potentially weakening its structural and functional efficacy (**Supplementary Table 1**). Consistent with these predictions, MutD also produced a greater increase in translational activity.

We used the MutD profile as a reference to extract additional candidate mutations from our dataset. Applying our criteria—90% structural identity, >2% increase in structural content, and lower MFE compared to WT—we also successfully retrieved the previously reported C→U mutation, which emerged as one of the top-ranking candidates (**Supplementary Figure 5**). This result validates our selection strategy and confirms that structural metrics derived from in silico mutagenesis can recapitulate known functional mutations.

Together, these results suggest that rational design of single-nucleotide substitutions, guided by structural metrics and thermodynamic models, is a feasible strategy for enhancing IRES activity.

### Identifying potential enhancing mutations in viral and non-viral IRES

Using the same selection framework established for c-MYC IRES, we screened viral and non-viral IRESs for candidate activity-enhancing mutations. Variants were selected if they satisfied three criteria: (i) structural identity ≥ 90% relative to the WT, ensuring preservation of the overall RNA fold; (ii) increased structural content (SC), measured as a higher percentage of base-paired nucleotides (SC Mut > SC WT) a feature often correlated with enhanced protein–RNA interactions and ITAF recruitment (74,94); and (iii) lower predicted minimum free energy (ΔG Mut < ΔG WT) indicating improved thermodynamic stability. We then mapped the resulting mutations onto each IRES sequence to analyze their positional distribution.

Interestingly, in viral IRES we identified a similar number of stability-promoting mutations, ∼10% of the overall mutants. Specifically, we found in PV the highest number of stability-promoting mutations (10.8%), followed by CrPV (10.5%), HCV (9.7%), and FMDV (9.5%). These mutations cluster mainly in the 5′ segment of the IRES—particularly in FMDV, CrPV, and PV, including several variants that strongly stabilize the RNA structure (**Figure 6A**). A secondary hotspot for beneficial mutation appears in the mid-sequence of the IRESs, after a valley with fewer beneficial mutations (250-300 nt for FMDV; 150-200 nt for PV). By contrast, HCV displays a more homogenous beneficial mutation landscape. The HCV IRES adopts a compact architecture in which individual domains participate in successive stages of pre-initiation-complex assembly, distributing functional contacts along the full RNA (9,95). In particular domains III and IV form most of the interaction surface for both the 40S subunit and eIF3 (41,43,59). Consequently, stabilizing mutations might be beneficial across the whole sequence. We identify a higher number of beneficial mutations in the 5′ (first 50 nt) compared to the 3′ (last 50 nt) for all the viral IRES sequences, reaching a 3-times higher density of mutations in the 5′ of PV. Overall, in all four cases, the 3’ sites of the IRESs are generally lacking beneficial mutations (**Figure 6A** and **Supplementary Figure 6**). Thus, few single-nucleotide substitutions in the IRES 3′ regions meet all three “beneficial” criteria, likely because the sequence immediately upstream of the AUG contains fewer modifiable structural motifs leaving little room for further stabilisation. This constraint may be especially relevant for type I and type II IRESs, which act as flexible scaffolds. In FMDV (type II), ribosomes bind the first AUG, and translation from the second AUG follows a scanning step from that site (96). In poliovirus (type I), the ribosome is initially recruited to an upstream AUG and then scans through domain VI to reach the start codon (97,98). Such scanning-dependent initiation likely favors reduced structural content at the 3′ boundary of the IRES. It was found that strong eukaryotic IRESs tend to have weak secondary structure immediately upstream of the AUG, consistent with a benefit of local de-structuring in that region (99).

**Figure 6.**
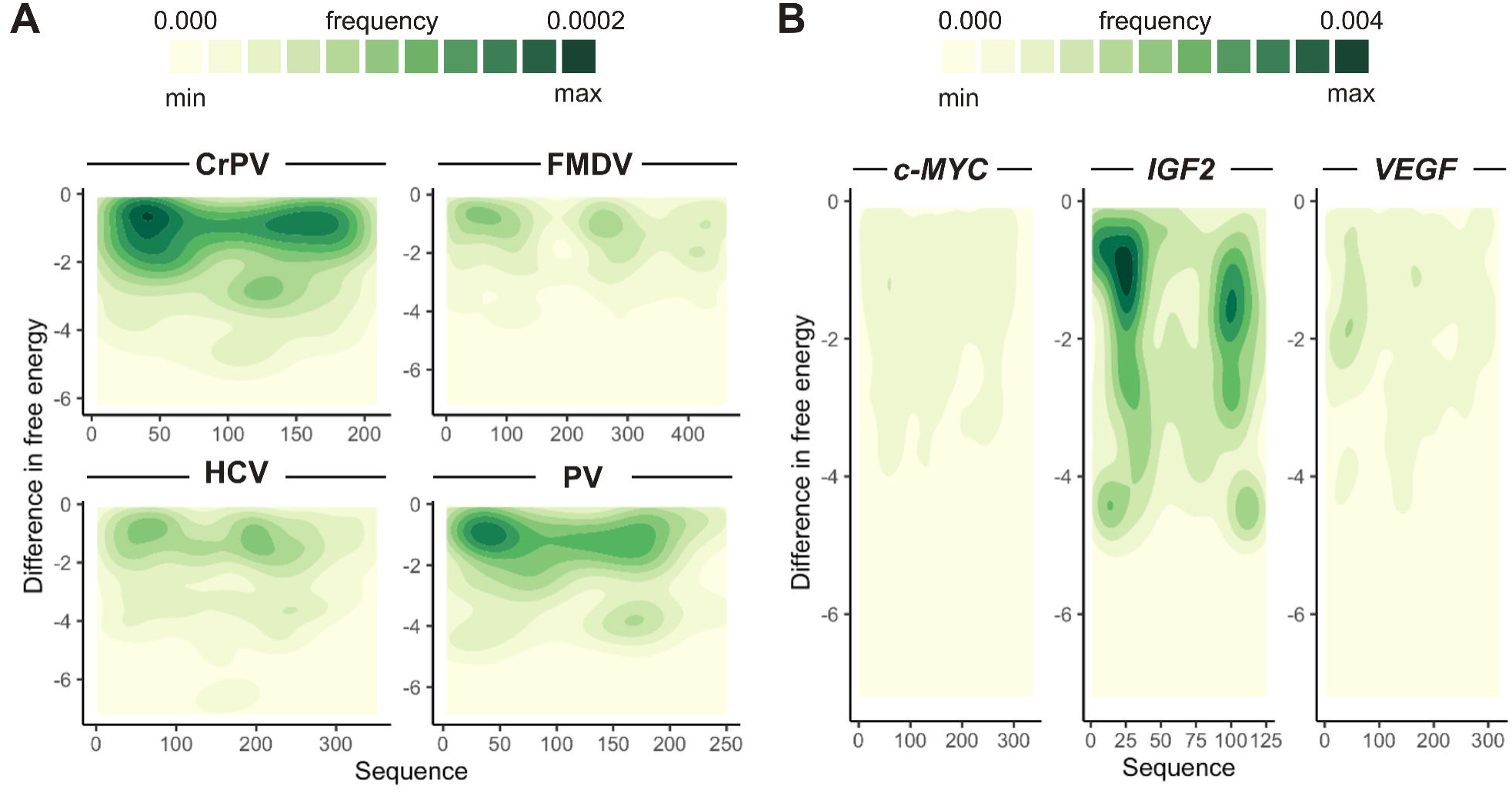
Distribution of IRES beneficial mutations. Frequency of mutations with structural identity ≥ 90% relative to the WT increased structural content viral IRESs (**A**) and non-viral IRESs (**B**). The frequency of mutations goes from low (light yellow) to high (dark green).

Non-viral IRES tend to have a slightly less homogeneous number of beneficial mutations compared to viral IRES, with the highest number of mutations present in *VEGF* (13.1%). On the contrary, *IGF2* shows the lowest amount of beneficial mutations (8.1%). c-*MYC* IRES has a similar pattern compared to viral IRES (10.8%). Although *IGF2* yields the lowest overall hit rate, it displays the highest density of strongly stabilising mutations both in the 5′ and 3′ of its sequence (**Figure 6B**). Overall, these observations highlight the element-specific nature of non-viral IRES architecture and argue for tailored, context-aware optimisation strategies guided by mutational and interaction profiles, rather than a single generalizable rule set. Non-viral IRESs are highly heterogeneous, differing in sequence, structural organisation, and reliance on auxiliary factors, which precludes a single, generalisable design rule; our mutational profiling underscores this diversity. Moreover, increased structural stability is not universally advantageous. In specific cases, local de-structuring appears to facilitate factor access and enhance activity—for example, ITAF binding to the *APAF1* IRES promotes local unfolding and increases initiation efficiency (9). Increase in G-quadruplex stabilization in the VEGF IRES inhibits the IRES activity (25).

### Mutations increasing the binding with proteins

The mutations identified here are putatively beneficial under our working criteria—preserving the global fold while increasing base pairing and lowering free energy. However, we do not exclude that mutations reducing local structural content may also be beneficial in certain contexts. Larger structural rearrangements might likewise enhance predicted ITAF/initiation-factor binding, yet beyond an unknown threshold the element may cease to function as an IRES. Experimental validation is therefore essential; our “beneficial” set should be viewed as high-confidence, structure-preserving candidates rather than an exhaustive catalogue of activity-enhancing variants. *In vitro* studies show that IRES elements are typically weak on their own and require the right ITAF to achieve efficient translation (86). ITAFs remodel IRES architecture to facilitate ribosome engagement and can also act as adaptors linking the RNA to the ribosome and initiation factors. Because IRES activity depends on recruiting the translation machinery, we evaluated it *in silico* by scoring predicted interactions with ribosomal components and initiation factors, *i.e.* recruitment potential.

First, we investigated whether the selected stabilizing mutations enhance binding to known ITAFs. For example, PV has four characterized ITAFs—PCBP2, GARS, La autoantigen, and RACK1 (74). We used human proteins to approximate the binding also for non-human IRES sequences based on the high sequence conservation of ITAFs among different species, with an average of 76 species with >90% conservation for 22 ITAFs (**Supplementary Table 2**). For example, RACK1 has 100% conservation with other 165 species, according to uniprotKB data (see **Materials and Methods**).

We examined which of the predicted beneficial mutations increase the binding affinity to these proteins. We identified mutations mildly (*cat*RAPID score MT - *cat*RAPID score WT > 5) and highly (*cat*RAPID score MT - *cat*RAPID score WT > 10) increasing the binding with the ITAFs. These mutations predominantly localize at the 5′ and 3′ ends of the IRES, as well as around the ∼175–200 nt region. Moreover, we also identified mutations able to increase 5-fold the binding with the ITAFs, mainly localised in the terminal IRES regions. Interestingly, we pinpointed 12 mutants increasing by 10-fold the binding with the ITAF, specifically with PCBP2. PCBP2 binds and stabilizes the apical region of stem–loop IV within the IRES, playing an essential role in facilitating ribosome docking (74,100). Therefore, enhancing the interaction between this protein and the PV IRES could promote greater structural stabilization and facilitate more efficient ribosome recruitment.

Next, we examined literature-supported ITAFs for the FMDV and *c-MYC* IRESs. For FMDV, we predicted the interaction with PTBP1, ITAF45, GEMIN5, SAM68, RAB1B, DDX3, NCL, G3BP1, HNRP, DDX21, DDX23, coherently with what is reported in the literature (101). For *c-MYC*, we evaluated the ITAFs hnRNPC1/2, hnRNPE1/2, hnRNPK, hnRNPL, DAP5 (37,38,102). Across single-nucleotide variants, *c-MYC* shows the largest fraction of structurally beneficial mutations—those that increase structural identity—that also enhance ITAF binding (53%; **Supplementary Table 3**). By contrast, FMDV exhibits the strongest magnitude of binding gains with 27% of variants that improve binding yield ≥10-fold higher predicted interaction than WT (**Supplementary Table 3**). At the factor level, hnRNPK and DAP5 account for the highest share of ≥10-fold improvements in c-MYC (19% and 14%, respectively), whereas GEMIN5 and PTBP1 lead for FMDV (28% and 22%).

After focusing on the few known ITAFs, we decided to look at the beneficial mutations enhancing the IRES activity by binding with proteins associated with the translation machinery. First, we applied *cat*RAPID to determine whether the MutA and MutD in *c-MYC* IRES are predicted to strengthen interactions with ITAFs and/or eiF, that could ultimately enhance translation. Each mutant was screened against the human RBPome (>2,000 proteins; **Materials and Methods**), and their binding propensities were compared with those of the WT *c-MYC* IRES. We then ranked all the RBPs by the magnitude of their binding propensity compared to the WT and extracted the top 50 (2.5 %) for each mutant, the resulting lists were significantly enriched for translation-related factors (MutA enrichment p < 5.7×10^-8^; MutD p < 5.5×10^-4^) (**Supplementary Figure 7**). Increased affinity for translation-related RBPs therefore provides a useful *in silico* indicator of mutations that are likely to increase the “recruitment potential” and thereby augment IRES-mediated initiation, as observed for MutA and MutD. Therefore, after having identified structurally stabilizing mutants, we used *cat*RAPID to test whether their predicted affinity for the translation machinery exceeded that of the WT IRES. We compiled a reference set of 119 translation-related RBPs—excluding ribosomal proteins and translational repressors—to serve as our proxy for translation enhancement. We then computed the overall interaction propensities between the selected proteins and the IRES sequences harboring the selected structure-preserving, stability-promoting mutations previously defined. For each viral IRES, we compared *cat*RAPID interaction scores for mutant *versus* WT sequences and computed the percentage of the 119 translation-related proteins showing higher binding to the mutant (**Figure 7**, **Materials and Methods**). CrPV exhibited the smallest fraction of translation-related proteins with increased binding, underscoring its relative independence from protein cofactors. Our results indicate that IRES elements relying on larger regulatory protein networks give rise to mutants with the greatest predicted gains in translation-factor binding. This pattern reflects their distinct modes of action: the CrPV IRES, which operates almost autonomously through its compact and stable structure, shows minimal binding enhancement, whereas the poliovirus IRES—less structurally constrained yet reliant on multiple eIFs and ITAFs—exhibits a much stronger increase (103). Notably, the non-viral IRESs (*VEGF* and c-*MYC*) behave similarly to viral IRESs that depend on trans-acting factors, consistent with previous observations that non-viral IRESs are generally less structured than viral counterparts (38). In contrast, structure-preserving, stability-enhancing mutations in *IGF2* did not improve the recruitment of translation-related proteins. *IGF2* IRES may rely less on auxiliary RBPs/ITAFs and instead engage more directly with specific ribosomal proteins (*e.g*., specialized ribosomes marked by RPL10A/uL1) (104), which would limit gains from mutations that primarily enhance RBP binding. Overall, we found that the structural-stabilizing mutations tend to increase the interaction with translation-related proteins, highlighting novel strategies to optimize and design synthetic IRES constructs. However, we also noticed how the magnitude of this effect depends on the IRES type and on its functional dependence on co-factors.

**Figure 7.**
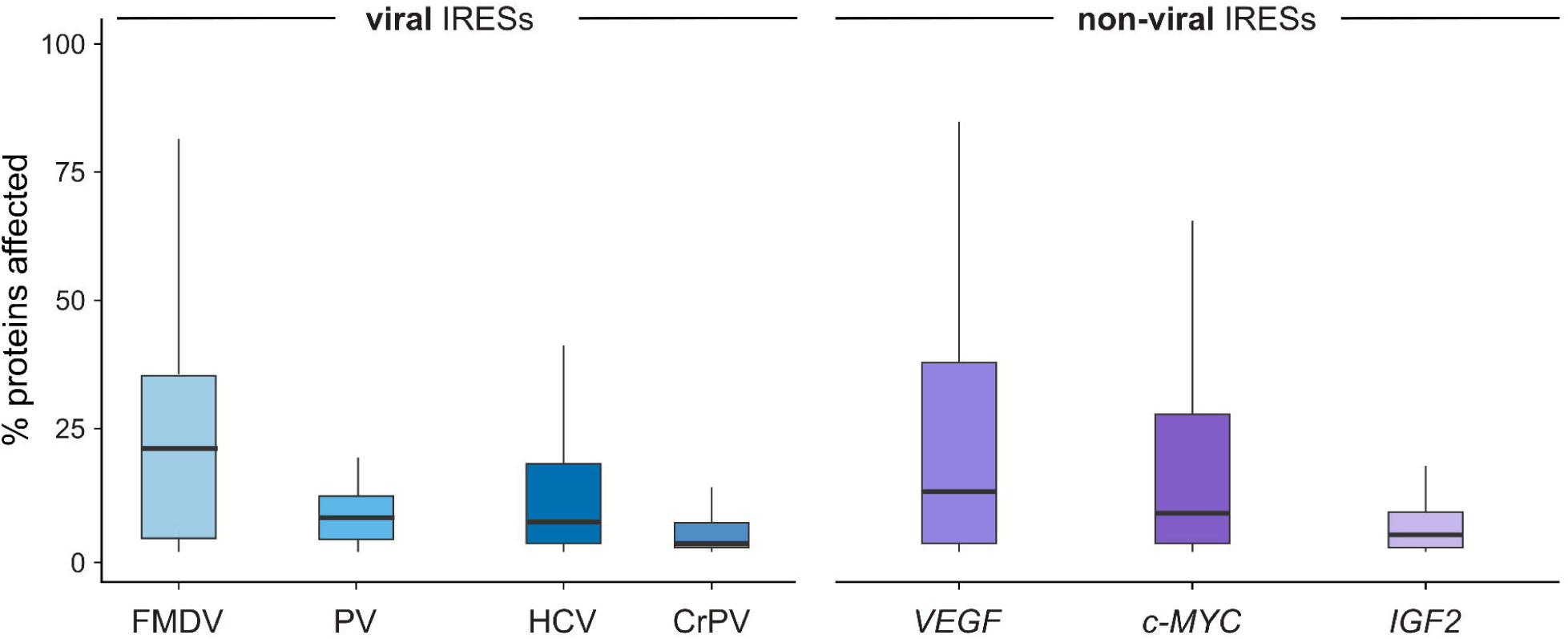
Translation enhancing effect of IRESs’ stability-promoting mutations. Percentage of RBPs with increased predicted binding to IRES variants carrying structure-preserving, stability-enhancing mutations, compared to the WT.

## Discussion

The 5′ UTRs of mRNAs, unconstrained by coding requirements, can host diverse RNA structures that regulate translation. Among these, IRESs enable cap-independent initiation and act as alternative regulatory modules. In RNA viruses, IRESs are essential for ensuring translation of the viral genome, thus ensuring viral fitness. On the other hand, non-viral IRESs play a key role during stress and in physiological contexts such as development (3,5,11,12).

In our work, we combined *in silico* mutational profiling, structural modeling, and protein–RNA interaction predictions to investigate how IRESs tolerate sequence variation. Mutational profiling has long been applied to understand and characterize functional RNAs such as ribozymes and riboswitches (47). This strategy captured both their structural robustness and their ability to engage regulatory proteins, providing insight into their mechanisms of action. Such an approach is particularly valuable given the low sequence conservation among IRESs, which complicates function prediction (14,38). Even well-characterized viral IRESs display distinct initiation modes, from factor-dependent scaffolds such as HCV and FMDV (43,105) to fully autonomous ribosome recruitment in CrPV (10).

Our analysis reveals different subsets of mutations having a different effect on the secondary structures, including single-nucleotide mutations completely altering the structure (non-synonymous structural mutations). We identified that IRES responses to non-synonymous mutations reflect their functional strategies: compact IRESs such as HCV are particularly sensitive in domains that recruit cofactors like eIF3 (60), while flexible elements like *IGF2* tolerate wider sequence variation (24). These patterns suggest different balances between rigid scaffolds, protein-assisted remodeling, and conformational plasticity.

This principle is especially relevant for non-viral IRESs. Once considered stress-specific, they are now known to act in normal developmental programs, including embryogenesis, as reported for *Hox* and *FGF1* (106,107). Recent studies have further expanded their repertoire (73). In addition, IRES-like activity has been identified in circRNAs, which may act in trans to promote translation of other transcripts (72,90).

A key advantage of our computational framework is its ability to prioritize regions for experimental validation by pinpointing structurally sensitive sites and protein-binding hotspots. For example, we identified single-nucleotide variants that stabilize the fold while enhancing predicted interactions with translation-related proteins. This mirrors known cases such as the C→U mutation in the c-MYC IRES, which promotes additional stem-loop formation and increases PTBP1 and YB-1 recruitment (19,65). Such examples demonstrate the potential of rationally designed synthetic IRESs. Considering that IRES efficiency can vary with tissue-specific ITAF expression (74), synthetic elements could be engineered to exploit these contexts, enabling selective translational control for biotechnology and therapeutics.

Finally, it is important to acknowledge the limitations of our approach. Since methods for 3D predictions of RNA structure such as Alphafold 3 are not ready for accurate high-throughput calculations (108,109), we have decided to use secondary structure predictions. Standard secondary structure calculations such as those carried out with the state-of-the-art RNAfold are optimized for speed and for handling multiple-sequence predictions, but this efficiency comes at the cost of a lower resolution. Additionally, these algorithms do not model pseudoknots, thereby reducing the computational complexity from NP-hard to polynomial time (110). Pseudoknots are often essential structural features of viral IRESs, such as the PKI domains of CrPV (63). Moreover, they do not account for G-quadruplexes that regulate activity in non-viral IRESs such as *VEGF* (25,27). The absence of these features may explain discrepancies between predicted mutational effects and experimentally observed activities. Integrating dedicated algorithms for pseudoknot and G4 modeling, without over-estimating the pseudoknot presence, will be crucial to refine mutational profiling and strengthen sequence–structure–function inference. We also focused on mutations that stabilise and promote the structure, however, we didn’t consider mutations decreasing the structural content while still stabilizing the energy.

## Conclusions

Our work provides a computational framework to analyze IRES elements through mutational profiling, structural modeling, and protein–RNA interaction analysis. By scanning sequence variants and predicting their impact on folding and binding, we could identify sites of structural fragility as well as mutations that enhance stability or favor protein engagement. This approach offers a means to prioritize IRES regions for experimental validation and to dissect the molecular principles underlying cap-independent initiation (73).

A key finding is that IRESs do not conform to a single structural blueprint. Rather, they display heterogeneous solutions to achieve ribosome recruitment: some rely on rigid scaffolds, as in the CrPV IRES, where stable stem–loops organize factor binding (63), while others exploit flexible conformations remodeled by ITAFs, such as PV (74).

Common structural features include stem–loops, long-range base pairs, pseudoknots, and in some non-viral cases, RNA G-quadruplexes (25,27). However, no universal motif emerges, underscoring the diversity of IRES mechanisms compared to canonical cap-dependent initiation (5). This places IRESs in the broader landscape of 5′UTR regulatory elements alongside upstream ORFs and structured hairpins (111).

Protein interactions play a central role in defining IRES function. ITAFs such as PTBP1, La, and YB-1 stabilize otherwise labile conformations, remodel structures to expose ribosome entry sites, or act as bridges between the IRES and the translation machinery (19,74). The interplay between RNA architecture and protein assistance explains both the robustness and the tissue-specific variability of IRES activity, since ITAF expression is cell-type dependent (74).

Another important regulatory axis involves RNA chemical modifications, which add a functional layer not captured by sequence or structure alone. Modifications such as N6-methyladenosine (m^6^A), pseudouridine, and 2′-O-methylation reshape RNA folding, stabilize or destabilize structural elements, and modulate protein recruitment (112). More recent work indicates that such marks can also alter the dynamics of ribonucleoprotein complexes, fine-tuning translation efficiency under distinct cellular states (113). These findings reinforce the view that IRES activity is the product of multiple regulatory layers: primary sequence, higher-order architecture, protein interactions, and epitranscriptomic modifications.

Altogether, our study reinforces the concept that IRESs represent diverse, adaptable platforms for translation initiation, distinct from but integrated with other 5′ UTR regulatory mechanisms. By combining computational scanning of mutations with structural and interaction profiling, we outline principles that may inform both mechanistic dissection and the rational design of synthetic IRESs for biotechnology and therapeutic applications. Future advances will require integrating pseudoknot- and G-quadruplex–aware folding algorithms (63) with maps of RNA modifications and systematic protein–RNA interaction assays, thereby moving closer to a comprehensive understanding of IRES structure and function.

## Declarations

The authors declare that they do not have any conflict of interest.

## Abbreviation

(IRESs): Internal Ribosome Entry Sites
(ITAF): IRES trans-acting factor
(RBPs): RNA-binding proteins
(PV): Poliovirus
(FMDV): Footh-and-mouth disease virus
(CrPV): Cripavirus
(HCV): Hepatitis C virus

## Availability of data and materials

All our results are reported in the main paper and additional supporting files.

## Authors’ contributions

R.D.P. and G.G.T. designed and conceived the study. R.D.P., A.V., and L.B. performed the analysis and assembled the figures. R.D.P., A.V., L.B., and G.G.T wrote the manuscript. All the authors read and agreed with the content of the manuscript.

## Acknowledgements

The authors thank the members of Tartaglia’s group for their constructive discussions and valuable feedback. We also acknowledge the RNA Technologies Flagship @IIT for their insightful comments and support.

## Supplementary Figures

**Supplementary Figure 1.** Structural identity for each mutant obtained from randomly mutating a nucleotide in one of the other three.

**Supplementary Figure 2.** Difference in free energy between the WT and the MT with structural identity = 100%.

**Supplementary Figure 3.** Structural identity distributions for the viral (**A**) and non-viral (**B**) IRES sequences.

**Supplementary Figure 4.** Enrichment of known ITAFs in IRES interactions. A. ITAF enrichment calculated at different thresholds of RBPs ranked by *cat*RAPID score compared to 10 random interactomes. B. Distribution of *cat*RAPID binding propensity score of the 2064 RBPs binding to VEGF and IGF2 IRESs.

**Supplementary Figure 5.** Frequency of mutations of one nucleotide into another as beneficial mutations selected with filters on structural and free energy change.

**Supplementary Figure 6.** Presence of beneficial mutations in the 5’ (first 50 nt) and 3’ (last 50 nt) of IRES sequences.

**Supplementary Figure 7.** GO enrichment of the 50 highest-ranked RBPs by change in binding propensity (MutA/MutD sequences vs WT).

## Supplementary Tables

**Supplementary Table 1.** Characteristics of the MutA and D sequences identified in our dataset. Average structural content and free energy for the MT and the WT are reported in the Table.

**Supplementary Table 2.** Number of species for *22 ITAFs* with sequence conservation at 90-100% according to UniRef clustering.

**Supplementary Table 3.** Number of beneficial mutations increasing the binding with known ITAFs 2-fold (2x), 5-fold (5x), 10-fold (10x) compared to the WT. The % is related to the overall number of beneficial mutations.

