## Supplementary Materials for "Integrating RNA Structure and Protein Interactions to Uncover the Mechanisms of Viral and Cellular IRES Function"

### Supplementary Figures

**Supplementary Figure 1.** Structural identity for each mutant obtained from randomly mutating a nucleotide in one of the other three.

**Supplementary Figure 2.** Difference in free energy between the WT and the MT with structural identity = 100%.

**Supplementary Figure 3.** Structural identity distributions for the viral (A) and non-viral (B) IRES sequences.

**Supplementary Figure 4**. Enrichment of known ITAFs in IRES interactions. A. ITAF enrichment calculated at different thresholds of RBPs ranked by *cat*RAPID score compared to 10 random interactomes. B. Distribution of *cat*RAPID binding propensity score of the 2064 RBPs binding to VEGF and IGF2 IRESs.

**Supplementary Figure 5**. Frequency of mutations of one nucleotide into another as beneficial mutations selected with the above criteria.

**Supplementary Figure 6.** Presence of beneficial mutations in the 5’ (first 50 nt) and 3’ (last 50 nt) of IRES sequences.

**Supplementary Figure 7.** GO enrichment of the 50 highest-ranked RBPs by change in binding propensity (MutA/MutD sequences vs WT).

**Supplementary Figure 1**

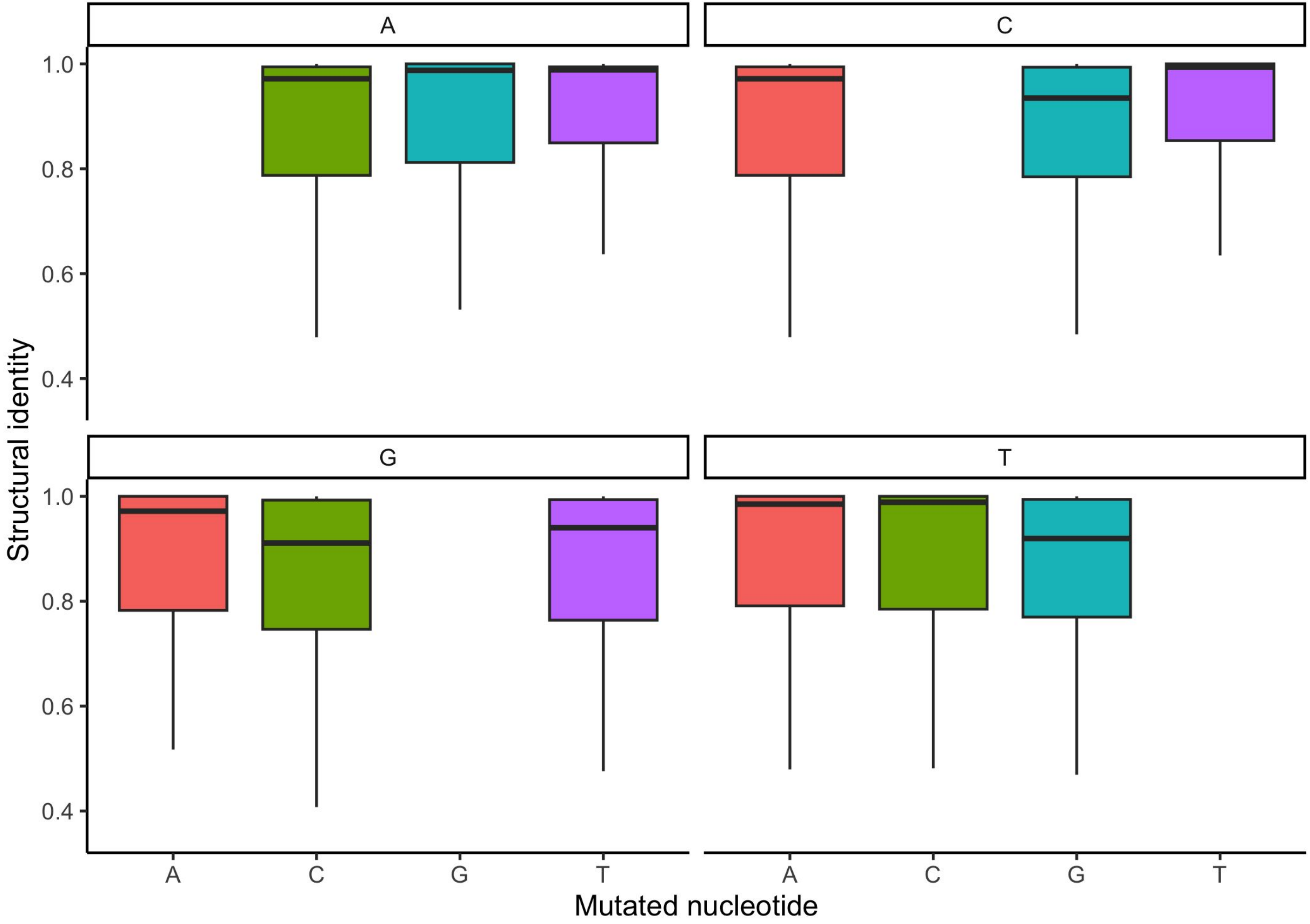

**Supplementary Figure 2**

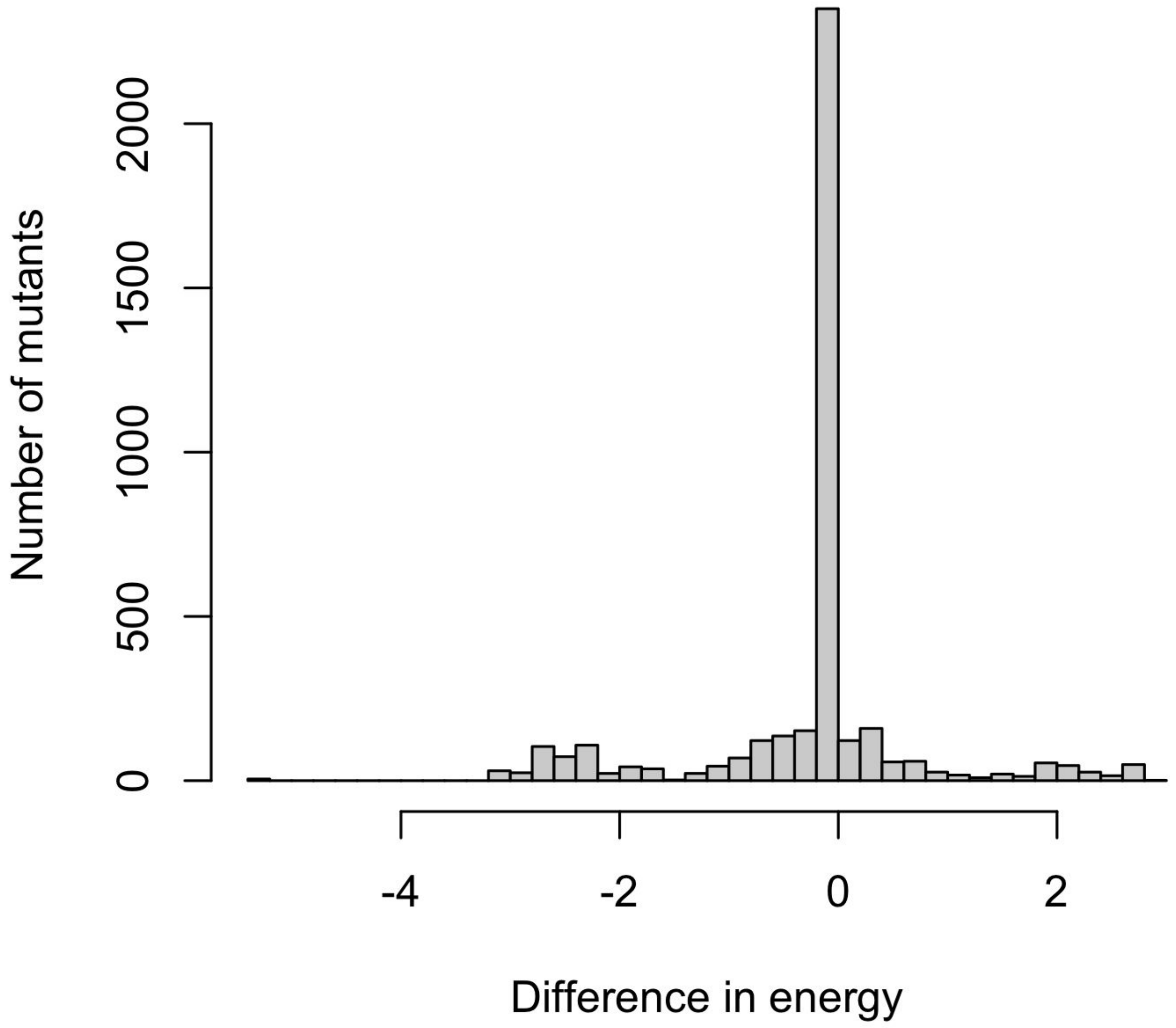

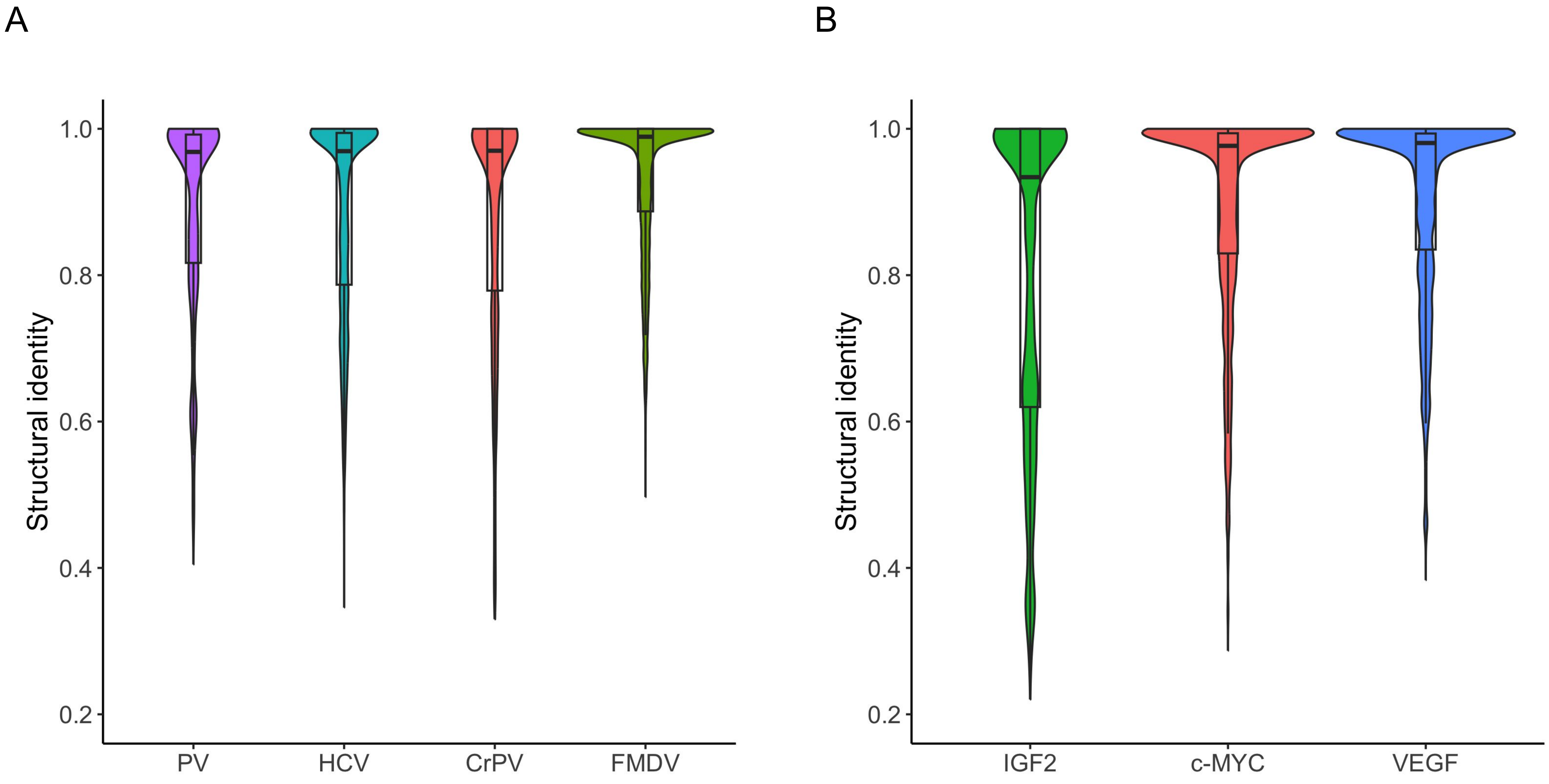


**Supplementary Figure 3**

Supplementary Figure 4

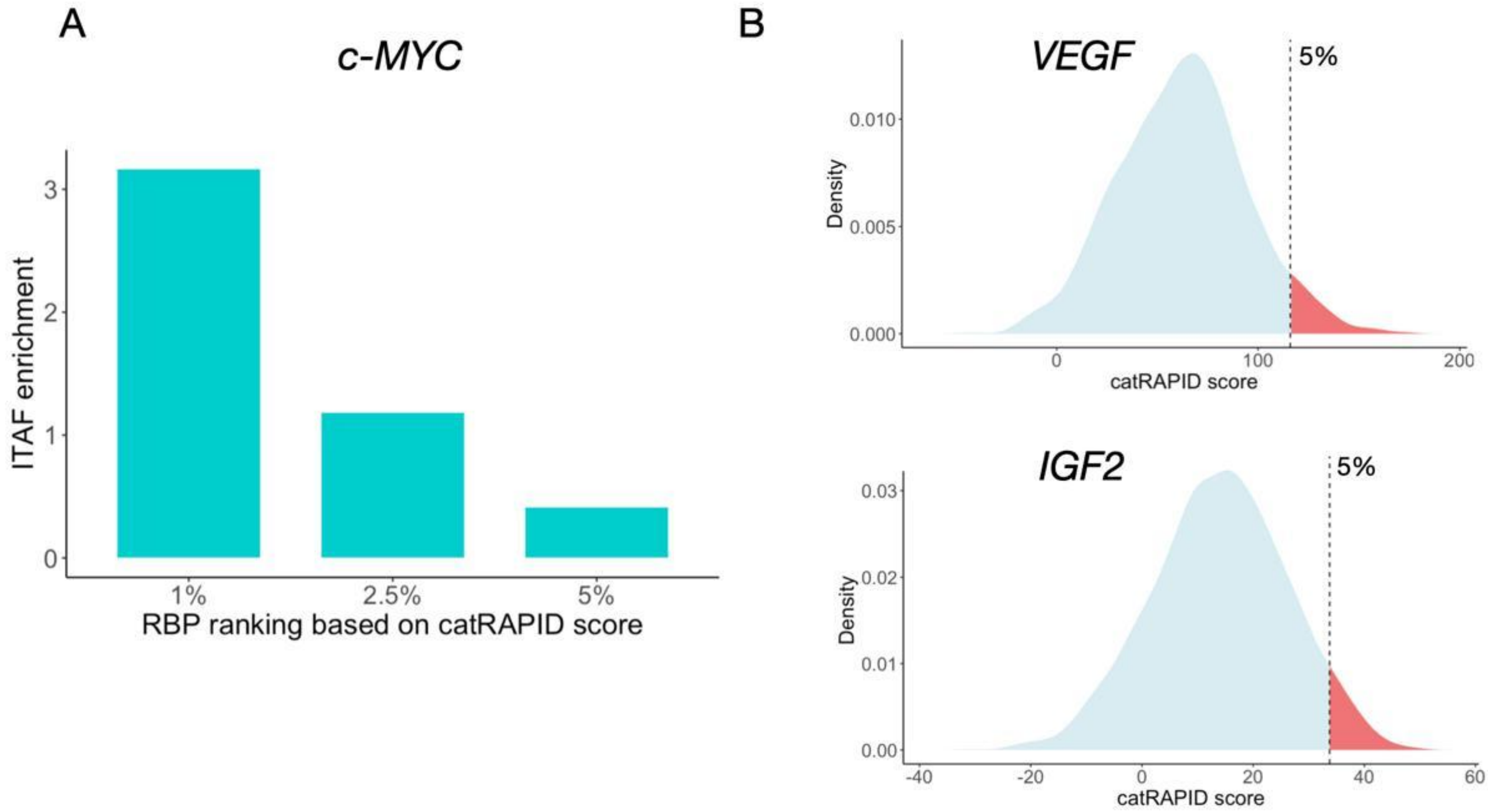

**Supplementary Figure 5**

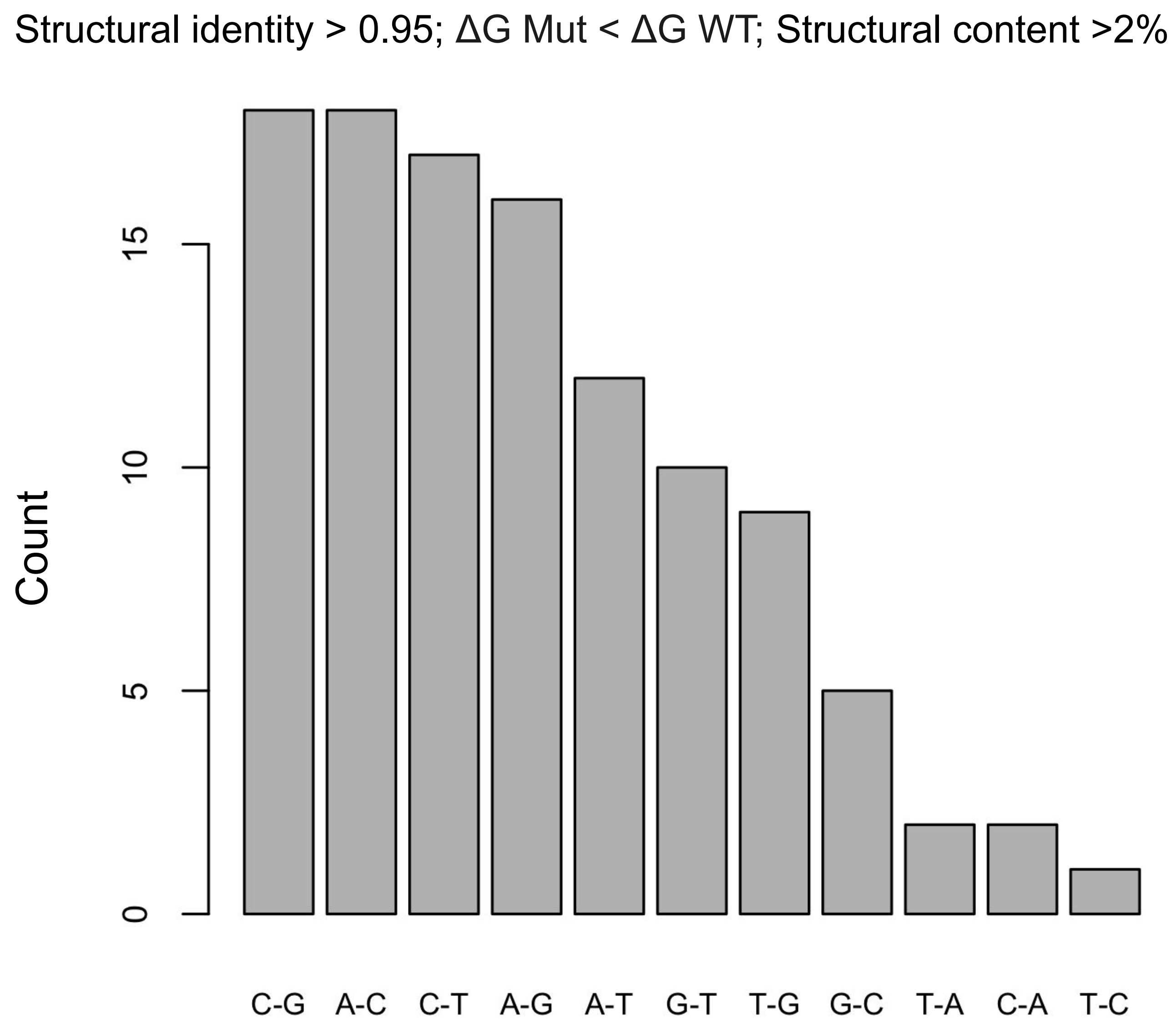

**Supplementary Figure 6**

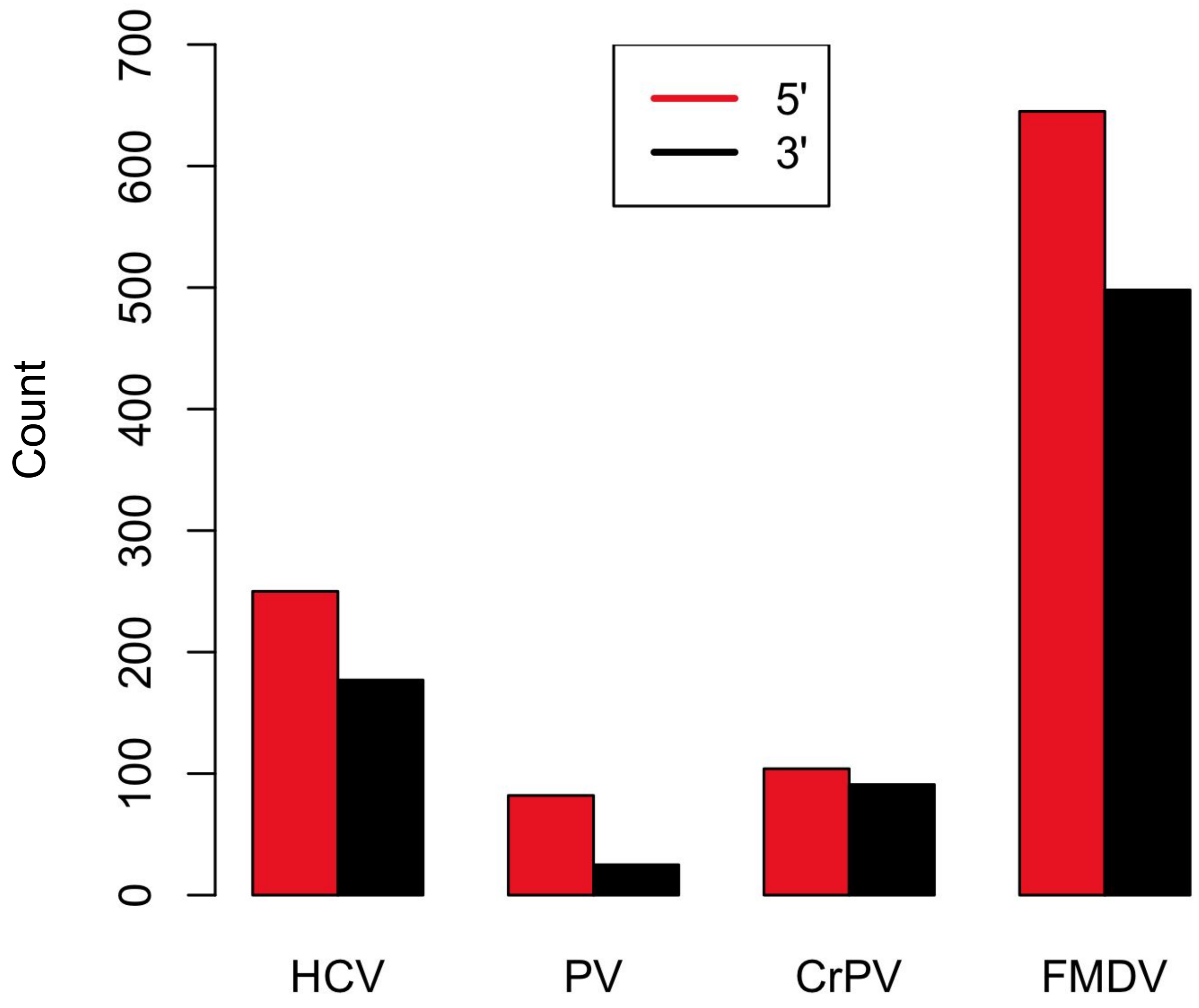

#### MutD

| Term Name | $p_{adj}$ (query_1) |
|---|---|
| RNA binding | $5.984\times10^{-17}$ |
| nucleic acid binding | $1.211\times10^{-10}$ |
| organic cyclic compound binding | $1.572\times10^{-9}$ |
| glutaminyl-tRNA synthase (glutamine-hydrolyzi... | $8.072\times10^{-3}$ |
| mRNA binding | $1.355\times10^{-2}$ |
| translation regulator activity | $1.402\times10^{-2}$ |
| catalytic activity, acting on RNA | $2.294\times10^{-2}$ |
| regulation of mRNA metabolic process | $1.486\times10^{-6}$ |
| regulation of mRNA stability | $1.596\times10^{-4}$ |
| regulation of RNA stability | $2.524\times10^{-4}$ |
| regulation of mRNA catabolic process | $3.026\times10^{-4}$ |
| mRNA metabolic process | $3.525\times10^{-4}$ |
| translation | $5.518\times10^{-4}$ |
| ribosome biogenesis | $6.544\times10^{-4}$ |
| mRNA catabolic process | $1.925\times10^{-3}$ |
| regulation of translation | $2.518\times10^{-3}$ |
| RNA catabolic process | $7.667\times10^{-3}$ |
| RNA metabolic process | $1.197\times10^{-2}$ |
| nucleic acid catabolic process | $1.328\times10^{-2}$ |
| gene expression | $1.513\times10^{-2}$ |
| negative regulation of translation | $1.557\times10^{-2}$ |
| glutaminyl-tRNAGln biosynthesis via transamid... | $2.421\times10^{-2}$ |
| nucleobase-containing compound metabolic pr... | $2.621\times10^{-2}$ |
| post-transcriptional regulation of gene express... | $2.912\times10^{-2}$ |
| organelle lumen | $1.948\times10^{-5}$ |
| membrane-enclosed lumen | $1.948\times10^{-5}$ |
| intracellular organelle lumen | $1.948\times10^{-5}$ |
| nucleoplasm | $1.217\times10^{-4}$ |
| intracellular membrane-bounded organelle | $9.743\times10^{-4}$ |
| nuclear lumen | $1.479\times10^{-3}$ |
| glutamyl-tRNA(Gln) amidotransferase complex | $3.366\times10^{-3}$ |
| nucleus | $4.273\times10^{-3}$ |
| intracellular organelle | $6.049\times10^{-3}$ |
| ribonucleoprotein granule | $7.011\times10^{-3}$ |
| membrane-bounded organelle | $1.622\times10^{-2}$ |
| P-body | $1.667\times10^{-2}$ |
| organelle | $4.503\times10^{-2}$ |

#### MutA

| Term Name | $p_{adj}$ (query_1) |
|---|---|
| RNA binding | $1.488\times10^{-12}$ |
| nucleic acid binding | $1.053\times10^{-9}$ |
| organic cyclic compound binding | $1.422\times10^{-8}$ |
| mRNA binding | $7.675\times10^{-6}$ |
| translation factor activity, RNA binding | $4.810\times10^{-5}$ |
| translation regulator activity, nucleic acid bindi... | $1.863\times10^{-4}$ |
| translation regulator activity | $7.333\times10^{-4}$ |
| catalytic activity, acting on RNA | $2.294\times10^{-2}$ |
| molecular condensate scaffold activity | $3.559\times10^{-2}$ |
| 3'-5'-RNA exonuclease activity | $4.598\times10^{-2}$ |
| translation | $5.715\times10^{-8}$ |
| mitochondrial gene expression | $5.406\times10^{-6}$ |
| mitochondrial translation | $2.831\times10^{-5}$ |
| biosynthetic process | $8.405\times10^{-4}$ |
| macromolecule biosynthetic process | $8.666\times10^{-4}$ |
| gene expression | $9.890\times10^{-4}$ |
| ribosome biogenesis | $1.138\times10^{-2}$ |
| translational initiation | $1.606\times10^{-2}$ |
| regulation of translation | $3.624\times10^{-2}$ |
| rRNA metabolic process | $4.620\times10^{-2}$ |
| positive regulation of stress granule assembly | $4.928\times10^{-2}$ |
| organelle lumen | $9.303\times10^{-6}$ |
| intracellular organelle lumen | $9.303\times10^{-6}$ |
| membrane-enclosed lumen | $9.303\times10^{-6}$ |
| intracellular anatomical structure | $8.019\times10^{-3}$ |
| cytoplasmic stress granule | $1.201\times10^{-2}$ |
| mitochondrial matrix | $2.291\times10^{-2}$ |

**Supplementary Figure 7**
